# Revisiting structural organization of proteins at high temperature from network perspective

**DOI:** 10.1101/2023.07.24.550270

**Authors:** Suman Hait, Sudip Kundu

## Abstract

Interactions between distantly placed amino acids in the primary chain (long-range) play a very crucial role in the formation and stabilization of the tertiary structure of a protein, while interactions between closely placed amino acids in the primary chain (short-range) mostly stabilize the secondary structures. Every protein needs to maintain marginal stability in order to perform its physiological functions in its native environment. The requirements for this stability in mesophilic and thermophilic proteins are different. Thermophilic proteins need to form more interactions as well as more stable interactions to survive in the extreme environment, they live in. Here, we aim to find out how the interacting amino acids in three-dimensional space are positioned in the primary chains in thermophilic and mesophilic. How does this arrangement help thermophiles to maintain their structural integrity at high temperatures? Working on a dataset of 1560 orthologous pairs we perceive that thermophiles are not only enriched with long-range interactions, they feature bigger connected clusters and higher network densities compared to their mesophilic orthologs, at higher interaction strengths between the amino acids. Moreover, we have observed the enrichment of different types of interactions at different secondary structural regions.

## INTRODUCTION

Folding of proteins to their characteristic three dimensional (3D) structures is an important aspect for every living organisms. The underlying mechanisms of protein folding and stability is yet to be completely understood.^1,2^ Various theoretical studies, which include simulations, statistical comparison and machine learning, have been performed on available protein structures to understand the mechanism of protein stability and to propose a guideline for enhancing it.^3–5^ Beside there are experimental works that try to engineer thermostable enzymes incorporating the knowledge from preceding theoretical end experimental studies.^6,7^ Even after rigorous research over several decades, researchers are unable to concur with any universal adaptation mechanism of thermostability, rather, it has appeared to be an interplay of several molecular features, encoded underneath the 3D structures.^8–10^ During protein folding, amino acids make several contacts among themselves, which play crucial role in determining the native 3D structure of the protein. These contacts can be represented as an amino acid contact network where the nodes are denoted by amino acids and an edge between two nodes is considered if the respective amino acids occur within certain distance cutoff in 3D.^11^ If two interacting nodes, *i* and *j*, in 3D space, are separated by more than 10 amino acids in the primary chain (|*i* − *j*| > 10), they form long range interaction network (LRN).^11^ On the other hand, nodes within 10 amino acid separation in the primary chain form short range interaction network (SRN, where, |*i* − *j*| > 3 and |*i* − *j*| ≤ 10, a lower cutoff of 3 amino acid separation in the primary chain is taken to minimize the effect of secondary structure in the 3D contact network).^11^ This network or graph-theoretical approach is widely used in deciphering several inherent features of residue contacts in various studies. It has also been used to compare several network properties of contact networks and their sub-networks of thermophilic and mesophilic proteins.^4^

In our last two studies we have shown that thermophilic proteins feature more salt-bridges, disulfide bridges, cation-π, π-π and Coulombic interactions compared to their orthologous mesophilic counterparts, but how these interacting amino acids are positioned in the primary chain and whether they have any association with thermostability, were not tested.^9,10^ Two interacting amino acids in the three dimension, when close to one another in the primary chain, would stabilize local structures, while if distantly placed in primary chain, would stabilize the tertiary structure and have a larger impact on global stability.^11,12^ Previously Sengupta and Kundu, comparing 12 meso-thermo orthologous protein pairs, showed that the length-normalized size of largest connected component (LCC) of long-range interaction network (LRN) is bigger in thermophilic than mesophilic at higher interaction strength cutoff, while short-range interaction network (SRN) exhibits no such trend.^11^ In the current study, with a large dataset of 1560 orthologous protein pairs of different folds and functions,^9^ we try to explore-whether the above mentioned observations are valid irrespective of wide variety of size, fold, class and function of the proteins and to further analyze the association of network density and salt bridge, π-π, cation-π and electrostatic interactions etc, at long and short range, in thermostability. We observe that the number of different interactions in orthologous thermo-meso protein pairs, differs more significantly in long range than short range. We observe a pattern of thermopholic LRN having larger connected components and higher density than their mesophilic counterparts for nearly 95% and 85% of the total 1560 orthologous pairs, at higher interaction cutoff (*I*_*min*_, discussed in materials and methods section). Whereas, no significant difference is observed in these parameters in SRN. While number of interactions in thermophilic SRN is higher than their mesophilic orthologs, no statistical significance is observed in LRN. We further analyze, how these interactions are distributed among the secondary structural elements in a protein to find out a possible explanation for two opposing pattern in Coulombic and van der Waals interaction networks. This study, in its limited scope, with its simple experimental framework, is able to capture several important features of residue contact network and their association with themostability.

## MATERIALS AND METHODS

### Data collection

Comparison of two thermo-meso orthologous proteins based on only their evolutionary ancestry, without considering their global topologies, structures and functions often lead to inconsistent and contradictory outcomes.^9,10^ Therefore, in the current study we have used a very carefully curated dataset that we have used in our two previously published studies.^9,10^ This dataset contains the sequences and x-ray crystallographic structures (≤ 3.0 Å resolution, and ≥80% protein sequence coverage) of 1550 Meso-Thermo/Hyperthermo (M-T/HT) orthologous protein pairs which exhibit similar lengths, 3D topologies, ACO^13^ (ACO is calculated as the average separation of contacting amino acids in the primary chain), and same domain architectures. Here with these 1550 orthologous pair, we have combined 12 pairs from Sengupta and Kundu’s study.^11^ After removing the duplicates, this combined dataset contains a total of 1560 Meso-Thermo orthologous proteins pairs (**Data S1**). Our dataset includes proteins from 82 bacteria and 14 archaea distributed in mesophilic, thermophilic and hyperthermophilic groups based on their optimal growth temperature. The lengths of the proteins vary in the range from 50 to over 1000 amino acids. The ACO of the proteins vary in a wide range of 13 to 107.^9^ The orthologous pairs contain nearly 300 different Pfam^14^ domains. So, our input dataset is free from any kind of biasness.

### Finding salt-bridge, cation-π, π-π and di-sulfide bridge

Primary structure of protein is a one dimensional arrangement of twenty different amino acids, connected through peptide bonds. During folding, distant parts of this primary chain come closer and form different kinds of molecular contacts such as salt-bridge, di-sulfide bonds, cation-π, π-π contacts, van der Waals interaction etc.^9^ Salt bridges are strong Columbic interactions between side chains of oppositely charged residues approximately ≤ 5 Å apart.^15^ Oxidization of the sulfhydryl (SH) groups of cystine residues spaced within approximately 2.3 Å in protein structures form disulfide bonds.^16^ A π-π interaction is created when the distance between the centroids of two aromatic side chain are placed between 4.5 to 7.0 Å.^17^ Cation-π interaction is formed when the positively charged side chains of lysine and arginine and the aromatic rings of aromatic amino acids fall within a distance of approximately 4 Å.^18^

### Finding out van der Waals and electrostatic interactions

Van der Waals interaction between two residues is considered when their atoms comes within a cut-off distance of their combined van der Waals radii + 0.5 Å.^19^ All these interactions have been found to subsequently enhance the stability of a protein in various literatures. Our own in-house python scripts are used to identify these interactions.

Electrostatic interaction between two charged amino acids is considered when at least one ion from each of the two different amino acids come within a distance cutoff ranging from 4 Å to 10 Å.^20^ For simplicity we have not calculated electrostatic interaction energy of the interactions. However, we can consider that higher the distance cutoff, lower is the electrostatic interaction energy for the same pair of ions.

### Calculating network strength

Protein contact networks are constructed considering the amino acids as nodes and van der Waals interaction among these nodes as edges. For the network formation, we have calculated the interaction strength of amino acid side chains given in by,^11,21^

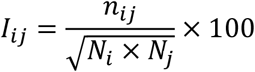

Where, *ni*_*j*_ is the number of distinct interacting pairs of side-chain atoms between the residues *i* and *j*, which come within distance d in 3D space (d = combined van der Waals radii of the atoms + 0.5 Å^19^). *N*_*i*_ and *N*_*j*_ are the normalization factors for the residues *i* and *j*, respectively. Normalization factor for the amio acid *i* is given as,^11,22^

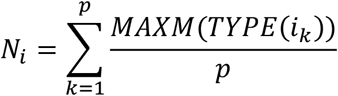

For a protein, *k*, all the side and main chain interactions of residue type *i* with its surrounding residues are calculated. *MAXM(TYPE(i*_*k*_)) is the maximum number of interaction pair by residue type *i* in protein *k*. The normalization factor for the residue type *i* (*N*_*i*_), is the average of *MAXM(TYPE(i*_*k*_))for total number of proteins in the study (*p*). The normalization factors consider the differences in different types of amino acids in terms of their sizes of the side chains and propensity of making contacts with other amino acid residues in protein structures.^11,22^ Using the same methodology, the interaction strengths for electrostatic interactions are also calculated for different distance cutoffs.

### Construction of long range network (LRN) and short range network (SRN) for van der Waals and Coulombic interaction

Once *I*_*ij*_ for all amino acid pairs are evaluated, the contact networks are formed, where an edge between nodes *i* and *j* is considered when *I*_*ij*_ is higher than a chosen cutoff value *I*_*min*_(*I*_*ij*_ >*I*_*min*_).^11^ This cutoff value is varied from 0% to 12% for van der Waalss interaction network and 0% to 25% for Coulombic interaction network in our study. Number of edges in the networks decreases with increasing cutoff as the number of nodes with higher number of interactions also decreases. The contact networks were then used to construct Long range interaction network (LRN, where, |*i* − *j*| > 10 in the primary chain) and Short range interaction network (SRN, where, |*i* − *j*| ≥ 3 and |*i* − *j*| ≤ 10 in the primary chain).^11^ The primary chain separation below three are not considered in the contact network as the immediate neighbors naturally feature large number of interacting atom pairs among them. Using the same methodology the LRNs and SRNs for all the orthologous pairs are calculated.

### Estimating the size of largest connected component (LCC) and network density (ND) for van der Waals interaction networks

For different cutoff values we have extracted the LCC for the LRNs and SRNs. A LCC is the largest group of connected nodes in a network that are reachable to each other directly or indirectly.^11,21,22^ Here, we have evaluated and compared the size and average network density (ND) of the LCC in LRNs and SRNs for each of the M-T/HT orthologous protein pairs. The ND of a network component is defined as the ratio of edges in the component over the maximum possible edges between the nodes in that component. For number of edges E and total number of nodes in a connected component N, ND is given by ND = 2E/N(N−1).^23^ Further the weighted average for the NDs of all connected components is calculated.

### Statistics and plot generation

PAST4.02 (PAleontological STatistics) software^24^ and our own in-house python scripts are used for the statistical analyses. All the plots are produced using OriginPro (ORIGINLAB, NORTHAMPTON, MA, USA) and Pyplot package of Python 2.7.

## RESULTS AND DISCUSSION

### LRN based on van der Waals interactions possess larger connected components even at higher interaction strength cutoffs

The role of van der Waals interactions in protein stability has been well documented in previous literatures. There are several studies of the network (created based on van der Waals interaction) parameters for M-T/HT orthologous proteins, but they cannot profoundly differentiate between meso and thermo, rather often provide contradictory outcome. Among several parameters the importance of the largest connected component (LCC) in network analysis has been well tested.^11,21,22^ LCC provides the information on nature and connectivity of a network and it undergoes a transition in its size as a function of the *I*_*min*_ cutoff for the whole contact network as well as for its different sub-network. Sengupta and Kundu showed that the difference in normalized size of LCC in thermo and meso is higher for long range network (LRN) than short-range network (SRN).^11^ Similar trend is observed in the normalized LCC vs *I*_*min*_for 1560 orthologous pairs in the current study. This pattern is not equally present in every orthologous pairs (**Supplementary Figure S1)**. Therefore, we take the mean value of normalized LCC at every *I*_*min*_cutoffs and have plotted the mean value in **Figure 1A**. The mean LCC size in SRN in meso and thermo do not possess significant difference in any of the *I*_*min*_cut-off while LRN in thermo holds a significantly higher (the *p* values of Mann Whitney-U test is given as gradient under the plot) value then LRN in meso even at higher *I*_*min*_ cutoffs (**Figure 1A**). The percentage increase in LCC size in thermo for SRN and LRN exhibit a significant difference in all the *I*_*min*_ cutoffs (**Figure 1B**) while the difference in their median value increases with increasing *I*_*min*_ cutoff. We then estimate the percentage of pairs for which the LCC size is bigger in thermophilic proteins than their mesophilic orthologs in all the higher *I*_*min*_ cutoffs than a selected cutoff *I*_*sel*_ . For that, we calculated the average LCC size for the orthologous proteins for all the *I*_*min*_ cutoffs, higher than *I*_*sel*_ . We also taken the percentage change of LCC size in thermophilic with respect to their mesophilic counterpart, which is denoted by Δ*LCC*_*SRN*_ for SRN andΔ*LCC* for LRN, where, Δ*LCC* can be expressed as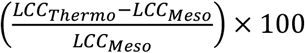. In **Figure 1C and 1D**, we have represented the percentage of orthologous pair, where LCC size is higher in thermophilic proteins than their mesophilic orthologs, with varying *I*_*sel*_ cutoff from 0% to 10% and Δ*LCC* cutoff from 0% to 20% in an incremental step of 5% for LRN and SRN respectively. At higher *I*_*sel*_ and higher Δ*LCC*_*SRN*_ cutoffs, the percentage of pairs with Δ*LCC*_*SRN*> 0_, drops below 30% (**Figure 1D**). On the other hand, for LRN, nearly 90% of the total pairs satisfy the trend at higher *I*_*sel*_ cutoff (*I*_*sel*_ ≥ 6%) even at Δ*LCC*_*LRN*_ ≥ 20% (**Figure 1C**). This clearly indicate that, at higher cutoff, the size of the largest connected component van der Waals interaction network in thermophilic proteins is bigger than their respective mesophilic orthologs, while the transition of LCC size for SRN is similar for thermophiles and mesophiles.

**Figure 1:**
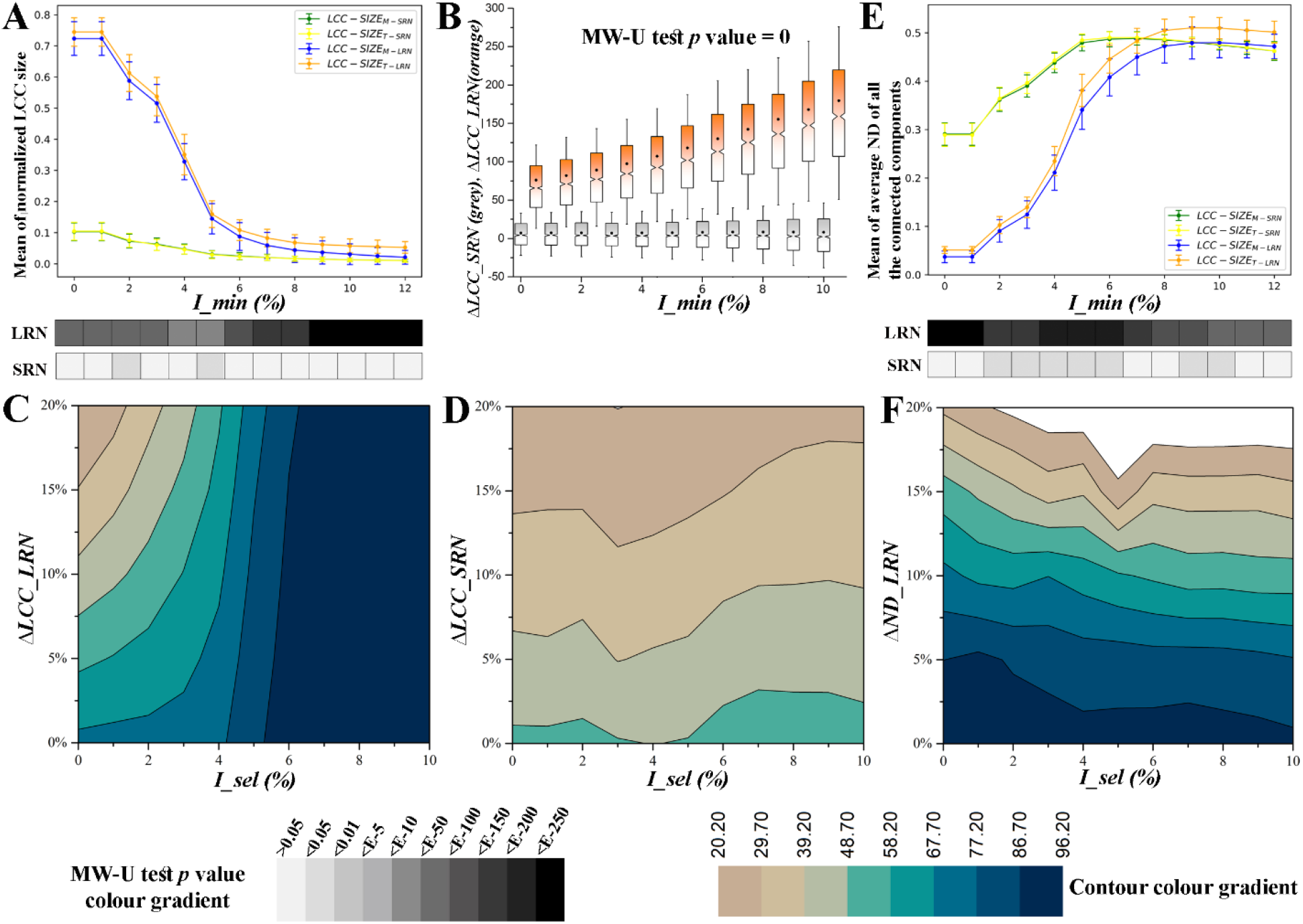
**(A)** Mean and standard deviation values of the normalized LCC sizes in 1560 mesophilic-thermophilic orthologous protein pairs are plotted against *I*_*min*_ cutoffs. The standard deviations are shown as error bars for the respective mean values. MW-U test *p* values of comparison between the LCC sizes in thermo and meso van der Waals interaction networks for LRN and SRN are shown below the x-axis. Darker blocks represent lower *p* values. **(B)** The percentages of mesophilic-thermophilic orthologous protein pairs for which normalized LCC size is bigger in thermophilic, are plotted as colored contour plots in the *I*_*sel*_ − Δ*LCC*_*LRN*_ space. **(C-D)** Δ*LCC*_*LRN*_ and Δ*LCC*_*SRN*_ plotted against *I*_*min*_ cutoffs. **(E)** Mean and standard deviations of the average ND values in 1560 mesophilic-thermophilic orthologous protein pairs are plotted against *I*_*min*_ cutoffs. The standard deviations are shown as error bars for the respective mean values. MW-U test *p* values of comparison between the NDs in thermo and meso Van der Waals interaction networks for LRN and SRN are shown below. Darker blocks represent lower *p* values. **(F)** The percentages of mesophilic-thermophilic orthologous protein pairs for which average ND values are bigger in thermophilic are plotted as colored contour plots in the *I*_*sel*_ − Δ*ND*_*LRN*_ space. (Please note that, for clarity of representation the standard deviation values are scaled down to 50% of their original values in **Figure 1A and 1E**)

Long-range interactions among amino acids in a protein contribute to the stability and integration of its tertiary structure. Moreover, the strength of interactions reflects the connectivity of different amino acids among themselves within a protein, influencing its packing and stability. The difference in the transition profiles of LCC in LRN between thermophilic and mesophilic orthologous proteins suggests that longrange interactions guide the enhanced stability of a protein’s tertiary structure in thermophiles. Whereas, larger long-range network components in thermophilic proteins indicate a need for more non-covalent interactions to bring together distant parts of a protein’s primary structure in three-dimensional space. At higher temperatures proteins naturally destabilize, one convenient way of preventing it could be creating higher number of interactions among the constituent amino acids.

### Denser LRN of van der Waal’s interaction network is a key feature of thermophilic proteins

A network is made of a set of connected components. The largest connected component alone is not enough for expressing protein stability from network point of view. All the connected components will also have some effect on the stability of a protein, based on how densely connected the nodes in all these components are. Therefore, we have calculated the average network density (ND) of the proteins for the components that contains at least three nodes. This average ND is calculated for different *I*_*min*_ cutoff varying from 0% to 12% and plotted using python matplotlib package (a sample of 4 pairs are provided in **Supplementary Figure S2**). The mean of average ND of all the connected components in a network for 1560 pairs is plotted against *I*_*min*_ cutoff in **Figure 1E** and **Supplementary Figure S2**. For SRN, the mean of average ND for thermo and meso appears to be very close to one another in every *I*_*min*_ cutoffs (**Supplementary Figure S2**). Whereas, for LRN, we observe significant difference among thermo and meso, in all *I*_*min*_ cutoffs. Also, the *p* value of significance gets higher in higher *I*_*min*_ cutoffs (*I*_*min*_ ≥ 5) (**Figure 1E**). We also estimated the percentage of pairs, for which the average ND value is consistently bigger in all *I*_*min*_ cutoffs higher than a selected cutoff, *I*_*sel*_ . Using a similar formula, we used for calculating Δ*LCC*, we also estimated Δ*ND*_*SRN*_ and Δ*ND*_*LRN*_ . The percentage of pairs with higher average ND value in thermophilic than their mesophilic orthologs, is represented in a contour plot for different Δ*ND* and different *I*_*sel*_ cutoffs for LRN and SRN in **Figure 1F** and **Supplementary Figure S3**, respectively. For Δ*ND*_*SRN*_ >0%, the percentage of pairs with higher average ND in thermophilic is roughly 40% for all *I*_*sel*_ cutoffs. Percentage of satisfying pairs drops gradually in all *I*_*sel*_ cutoffs with increasing Δ*ND*_*SRN*_ . For Δ*ND*_*SRN*_ ≥5%, the percentage of pair drops below 20%. On the other hand, in LRN, the percentage of pairs exhibiting higher ND value in thermophilic is near 90% at every *I*_*sel*_ cutoff for Δ*ND*_*LRN*> 0_ %. For Δ*ND*_*LRN*_ ≥5%, the percentage of satisfying pairs varies between 75-85% for different *I*_*sel*_ cutoff. Even for Δ*ND*_*LRN*_ ≥ 15%, percentage of pairs, satisfying the trend, occurs within 30-40%, for different *I*_*sel*_ cutoffs. This result implies that the amino acids (nodes) in LRN of thermophilic proteins are more densely connected than their mesophilic orthologs. Because of these densely connected nodes, thermophilic proteins can provide more resistance against thermal denaturation.

The network density (ND) of a protein refers to the level of connectivity among its residues. A higher ND indicates a greater number of interactions and a more densely connected protein structure. ND plays a crucial role in the stability of proteins. A densely connected network provides structural integrity and support, ensuring that the protein maintains its 3D shape. Therefore, changes in ND can be indicative of conformational changes or structural rearrangements in response to external stresses like temperature, salinity etc. Here the ND of SRN in both thermophilic and mesophilic orthologous proteins shown a similar transition profile over a wide range of *I*_*min*_ cut-offs, whereas ND in LRN shows significant difference. It clearly indicates that the need of thermal adaptation does not affect SRN but changes LRN for more interconnectedness between the nodes.

### Thermophilic and mesophilic proteins differ in SRN formed on the basis of Coulombic interactions

The LRN and SRN of the Coulombic interaction networks contain both attractive and repulsive interactions. But most of them do not form any cluster, rather occur as isolated interaction pairs in the network. So instead of estimating the LCC size, we took the number of pairs featuring higher interaction strength than a selected *I*_*min*_ cutoff. Then we take the net attractive−repulsive interaction count, *Net*− *coulombic*_*T*−*LRN*_ and *Net*− *coulombic*_*T*−*SRN*_ respectively for the LRN and SRN of thermophilic proteins. Similarly we estimate, *Net*− *coulombic*_*M*−*LRN*_ and *Net*− *coulombic*_*M*−*SRN*_ for orthologous mesophilic LRN and SRN respectively. Then we plot the mean value of these 4 parameters vs *I*_*min*_ for 1560 orthologous pairs for different distance cutoffs (**Figure 2A-D**). In contrary to van der Waals network, we observe that SRN possesses higher difference between thermo and meso and this trend weakens with increasing *I*_*min*_ value, when the distance cutoff for considering a Coulombic interaction is ≤10Å (**Figure 2A**). As we decrease the distance cutoff for Coulombic interaction the pattern does not change for the cutoff of ≤8Å and ≤6Å (**Figure 2B** and **2C**). But for the distance cutoff ≤4Å, we observe both of the LRN and SRN Coulombic interaction networks exhibit higher mean for both *Net*− *coulombic*_*T*−*LRN*_ and *Net*− *coulombic*_*T*−*SRN*_ . However, the mean value is higher for LRN for ≤4Å (**Figure 2D**), which means that the distant amino acids of the thermophilic proteins form higher number of stronger attractive (as the Coulombic interaction force increases with decreasing distance cutoff) Coulombic interaction than their mesophilic ortholgs do, although the effect of Coulombic interaction is limited to mostly the stabilization of the local structure.

**Figure 2:**
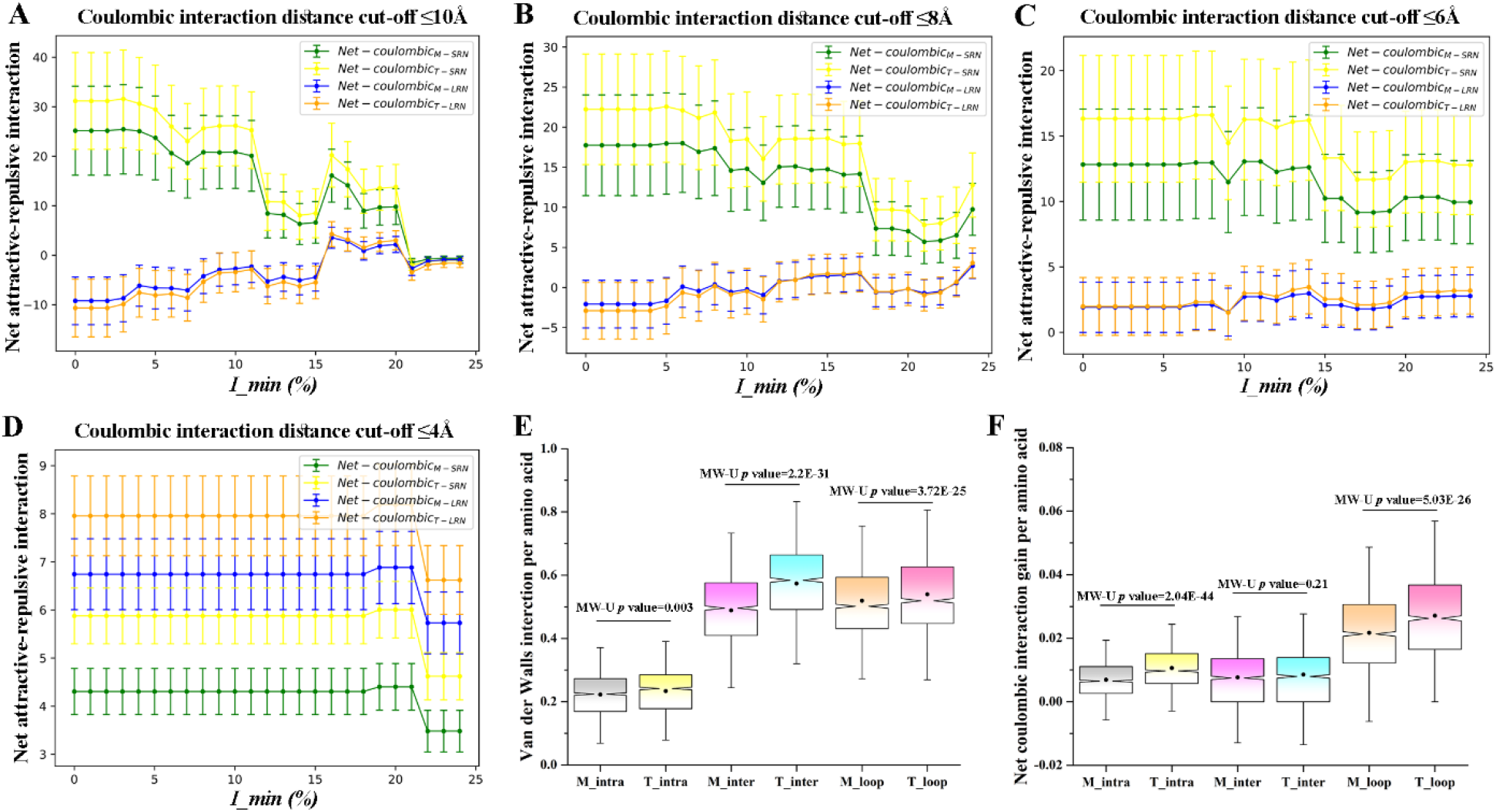
Figure **A-D** shows net attractive−repulsive Coulombic interactions vs *I*_*min*_ plots for different distance cutoffs. The standard deviations are shown as error bars for the respective mean values. **(E)** Distribution of van der Waals interactions in inter secondary structures, intra secondary structure and loop-linked groups. **(F)** Distribution of attractive−repulsive Coulombic interactions in inter secondary structures, intra secondary structure and loop-linked groups.

Coulombic interactions play an important role in the folding of proteins and also in maintaining their 3D structures.^25–27^ During protein folding, charged residues interact with each other and with the solvent, helping the protins to stabilize into specific conformations.^26,27^ These interactions help in the proper positioning of secondary structure elements as well as in the formation of tertiary structures.^26,27^ A Coulombic interaction occurs between charged residues that are spatially separated in the protein structure. Although the energetic contribution of Coulombic interactions are not as much as a salt-bridge, and it varies based on the distance and the charges of the ions, cumulatively these interactions can influence the stability of the protein by contributing to the overall electrostatic potential energy.^10^ Depending on the charges involved, Coulombic interactions can either stabilize or destabilize the protein structure. In the current study when we take the net attractive − repulsive Coulombic interaction count we consider all the interactions above a *I*_*min*_ cut-off contribute equally to the overall electrostatic potential energy. Therefore, an enhancement in this count indicates higher contribution in electrostatic potential energy. We have observed that thermophilic proteins feature higher number of salt-bridges in both LRN and SRN than their mesophilic orthologs (**Figure 3A**). Similarly, when we consider net electrostatic interactions of LRN and SRN we observe indistinguishable pattern for distance cutoff ≤4Å (close to cutoff of salt-bridge) (**Figure 2D**). However, when we consider higher distance cutoffs, the difference of net Coulombic interaction in SRNs of thermophilic and mesophilic is much higher compared to LRNs of the orthologs (**Figure 2A-C**). This indicates that Coulombic SRN aid more in achieving thermostability than Coulombic LRN.

**Figure 3:**
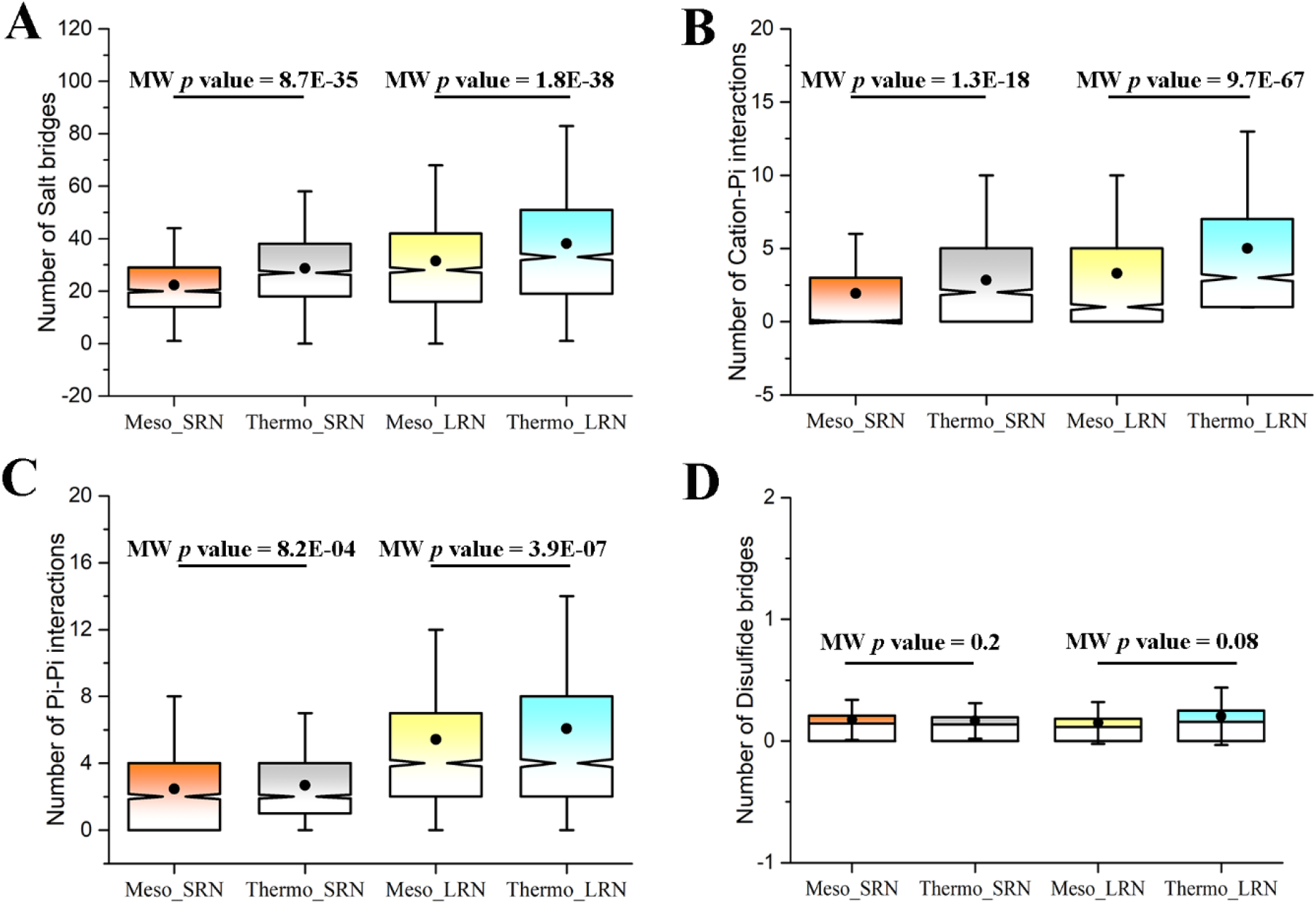
**A-D** Comparisons of the presence of salt-bridges (**A**), Cation-π (**B**), π-π (**C**) and Disulfide-bridges (**D**) in the LRN and SRN in mesophilic-thermophilic orthologous proteins.

### Stabilization of secondary structures and higher order structural organization

Van der Waals and Coulombic interactions are two of the five major interactions those play a key role in the folding and stability of proteins. The distribution of Coulombic and van der Waals’ interactions between different structural segments of proteins vary depending on the specific protein, its conformation, amino acid sequence, 3D structure, and the local environment of the protein. Thermophilic proteins are expected to have more van der Waals and Coulombic interactions,^9,10^ but at the same time it would be more interesting to find how these additional interactions in a thermophilic-ortholog are distributed within the protein structure.

The largest connected component (LCC) in van der Waals SRN and the number of net attractive−repulsive interaction in Coulombic SRN denote a strong local structural organization, whereas the same in LRN denote strong global structural organization of the contacting nodes. As we found that the thermophilic proteins feature higher number of net attractive−repulsive interaction in Coulombic SRN, we expect that Coulombic interaction will be over represented in intra secondary structural (intra SS) nodes, while van der Waals interactions are expected to be found in the inter secondary structural (inter SS) nodes. Distributing these interactions from both mesophilic and thermophilic orthologous proteins in three groups-i) intra SS, ii) inter SS and iii) loop-linked, we observe that thermophilic proteins are enriched with higher inter SS and loop-linked van der Waals interactions (**Figure 2E**) and intra SS and loop-linked Coulombic interactions (**Figure 2F**). As loop regions feature mostly polar and charged residues,^28^ they can be stabilized by forming both van der Waals and Coulombic interactions within and with the structured region.

### Salt-bridge, cation-π and π-π interactions are more abundant in thermophilic LRN than their orthologous mesophilic LRN

Folding of a protein to its native structure is achieved by clustering of hydrophobic patches followed by formation of several kind of interactions as distant regions of the proteins come closer. While interactions in SRN are involved in stabilization of mostly the secondary structures, interactions in LRN aid in holding two distant region of the proteins together. In our previous studies we have shown that thermophilic proteins feature higher number of salt-bridges, di-sulfide bridges, cation-π, π-π interactions as well as higher Coulombic energy gain coming from partially exposed charge reversal mutations but their association with thermostability at long and short range are yet to be explored. Here, an interaction is considered short range if the primary chain separation of the interacting residues are between three to ten and beyond ten it is considered as long range. So, we compare the number of these interaction (N(T) and N(M) are the number of a type of interaction in a thermophilic protein and it’s mesophilic counterpart, respectively) in short and long range for the orthologous pairs. In these comparisons, our null hypothesis H0 is N(T) > N(M) and the alternative hypothesis H1 is N(T) < N(M) for both long and short range. Salt bridges of both long and short range networks occur in higher number in thermophilic proteins compared to their mesophilic orthologs. The average number of salt bridges in short range are 22.3 and 28.7 in mesophilic and thermophilic, respectively. In long range the averages are 31.5 and 38.2 in mesophilic and thermophilic, respectively. The number of salt bridges in both mesophilic and thermophilic are higher in long range than their respective short range interaction. The *p* value of significance in Mann-Whitney U test between mesophilic and thermophilic in both long and short range exhibit similar values (8.7E−35 in short range and 1.8E−38 in long range, **Figure 3A**). The average numbers of cation-π interactions in mesophilic and thermophilic proteins are 1.9 and 2.8 in short range, respectively, whereas the same in long range are 3.3 and 5, respectively. In case of π-π interaction, these numbers are 2.4 and 2.7 in short range and 5.4 and 6.1 in long range in mesophilic and thermophilic, respectively. The *p* values of Mann-Whitney U test between thermophilic and mesophilic for both of these interactions show higher significance in long range than short range (**Figure 3B** and **3C**). Di-sulfide bridges are comparatively rarely found in the protein structures. When we distribute them according to long and short range, the number of di-sulfide bridges become even smaller. We could not found any significant difference between the Meso-Thermo/Hyperthermo (M-T/HT) orthologous proteins either in the short or long range (**Figure 3D**).

### Structural organization of thermophilic proteins depends on the long range interactions to a greater extent

Earlier studies have shown that, the normalized size of LCC undergoes a transition with increasing *I*_*min*_ cutoff for all the proteins and transition is sharper for SRN than LRN. Working on a dataset of twelve meso-thermo orthologous protein pairs, Sengupta and Kundu showed that the transition of normalized size of LCC in SRN is similar for both mesophilic and thermophilic proteins but the transition in LRN is sharper for mesophilic than thermophilic, which indicates the presences of larger interconnected long range interaction in thermophilic proteins.^11^ SRN is the network of interacting amino acids that are also closely positioned in the primary chain, hence provides local stability. On the other hand, LRN contain amino acids positioned far from each other in the primary chain but interacting in three dimension. So, LRN connects distant parts of a protein, establishes interactions between several secondary structures, thus stabilize the tertiary structure of a protein. We observe that with increasing *I*_*min*_ cutoff, the normalized LCC size gradually decreases in both SRN and LRN. To study this transition we estimated

*I*_*critical*_ value ^11^ (the *I*_*min*_ value at which the normalized size of LCC becomes half of the size it has at *I*_*min*_ cutoff = 0%) of all the networks for 1560 orthologous pairs. Lower the *I*_*critical*_ value for a plot, sharper is the transition, which mean the network loses its connections at a smaller *I*_*min*_ cutoff. The transitions in SRN in both mesophilic and thermophilic look similar (*I*_*critical*_ value falls with a range of 2%-2.25% for both meso and thermo SRN in 1560 pairs). The transition in LRN networks, are not as sharp as that in SRN. LRN networks drop down to half of its LCC size at higher *I*_*min*_ cutoff, i.e. higher *I*_*critical*_ value. However, the *I*_*critical*_ value for thermo_LRN (*Icritical*value is 4.0% to 4.5% for 1560 pairs) is even higher than meso_LRN (*I*_*critical*_ value is 3.5% to 4.0% for 1560 pairs). This means not the thermophilic proteins have higher number of interacting amino acid pairs also larger connected cluster of long range interactions at larger *I*_*min*_ cutoff. Besides our study also consider the effect of other connected components alongside the largest one. The average ND values of connected components is higher in thermophilic proteins than their mesophilic counterparts indicating densely connected nodes stabilizing thermophilic proteins even at higher *I*_*min*_ cutoff.

From previous literature we know that one of the key forces that stabilizes individual secondary structures is the formation of hydrogen bonds in a repetitive manner. Alpha helix and beta sheets are the two most stable polypeptide chains that have the highest hydrogen bonding potential.^29^ So, the higher number of Coulombic interaction within same secondary structure (intra SS in **Figure 2F**), is not the key force in stabilizing them, rather an associative one. Beside when we consider the distance cutoff for Coulombic interaction ≤4Å, higher number of interaction is found in thermophilic LRN, which means although a huge number of long distance (weak) Coulombic interaction gain is found in SRN, fewer but strongest Coulombic interaction is found in LRN network. Therefore, Coulombic interaction also have a key role in stabilizing global structure. Moreover, when different secondary structural elements are driven closer by hydrophobic interactions a large number of non-specific van der Waals interactions are formed between them. Beta sheet structure is also stabilized by formation of long range interaction between separate beta strands. Our result of over representation of van der Waals interactions in inter secondary structures and the loop-linked interactions supports the previous observations (**Figure 2E)**.

Our study also exhibit higher number of long range salt-bridge, cation-π and π-π interactions in thermophiles which may assist in stabilizing the tertiary structure at high temperature. Our analysis on a large, carefully curated dataset, reconfirms the outcomes of previous studies and found few more features of the association of thermostability with long range interactions, can encourage further exploration of the network aspect of thermostability.

## Author Contribution

S.K. and S.H. designed research; S.H. implemented computational methodologies and performed research; both analyzed data and wrote the paper.

## Acknowledgement

S.H. is supported by DBT-BINC SRF fellowship (Fellow Number: DBT-BINC/2017/CU/12).

## Abbreviations

LCC: largest connected components
LRN: long range network
SRN: short range network
*I*_*min*_: Interaction strength cutoff
ND: network density

