## Supplementary Figures for "Revisiting structural organization of proteins at high temperature from network perspective"

Hait and Kundu, 2023

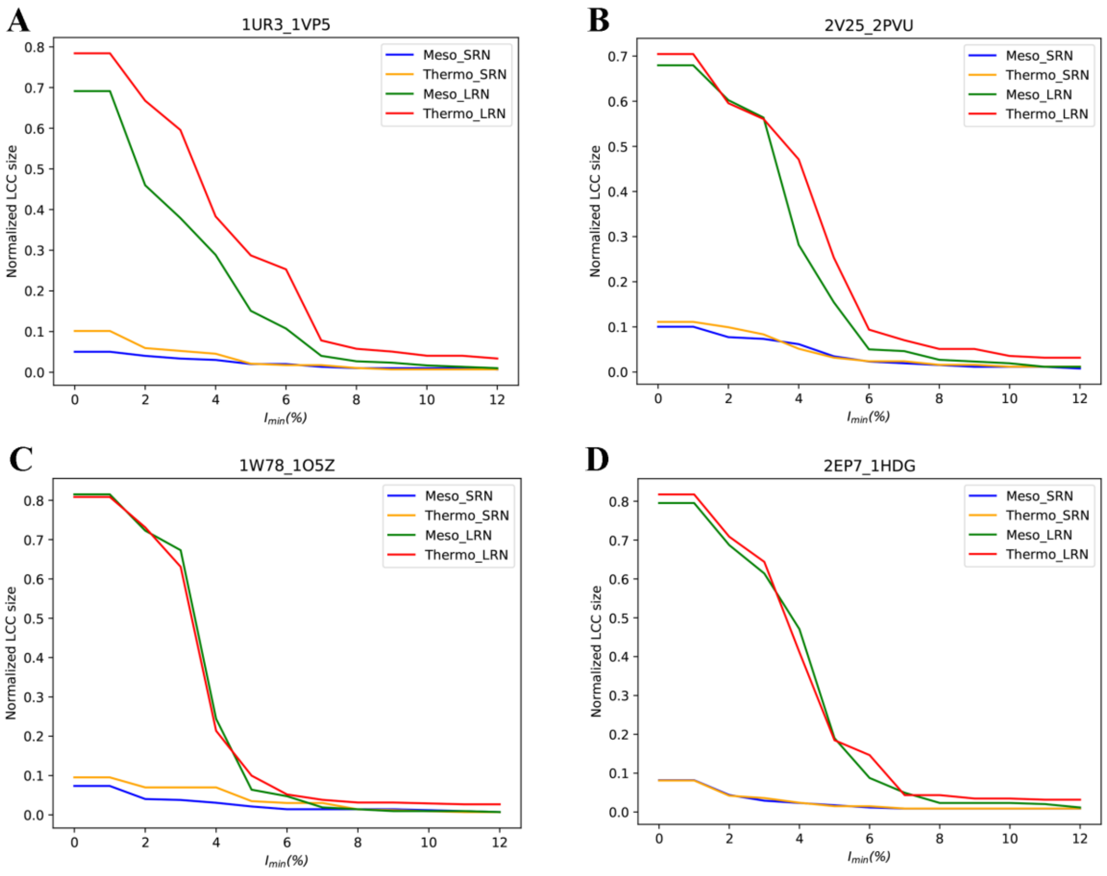


**Figure S1(A-D).** Sample Normalized LCC vs $I_{min}$ plot of four out of 1560 mesophilic-thermophilic orthologous pairs. The mean plot is provided in the main text.

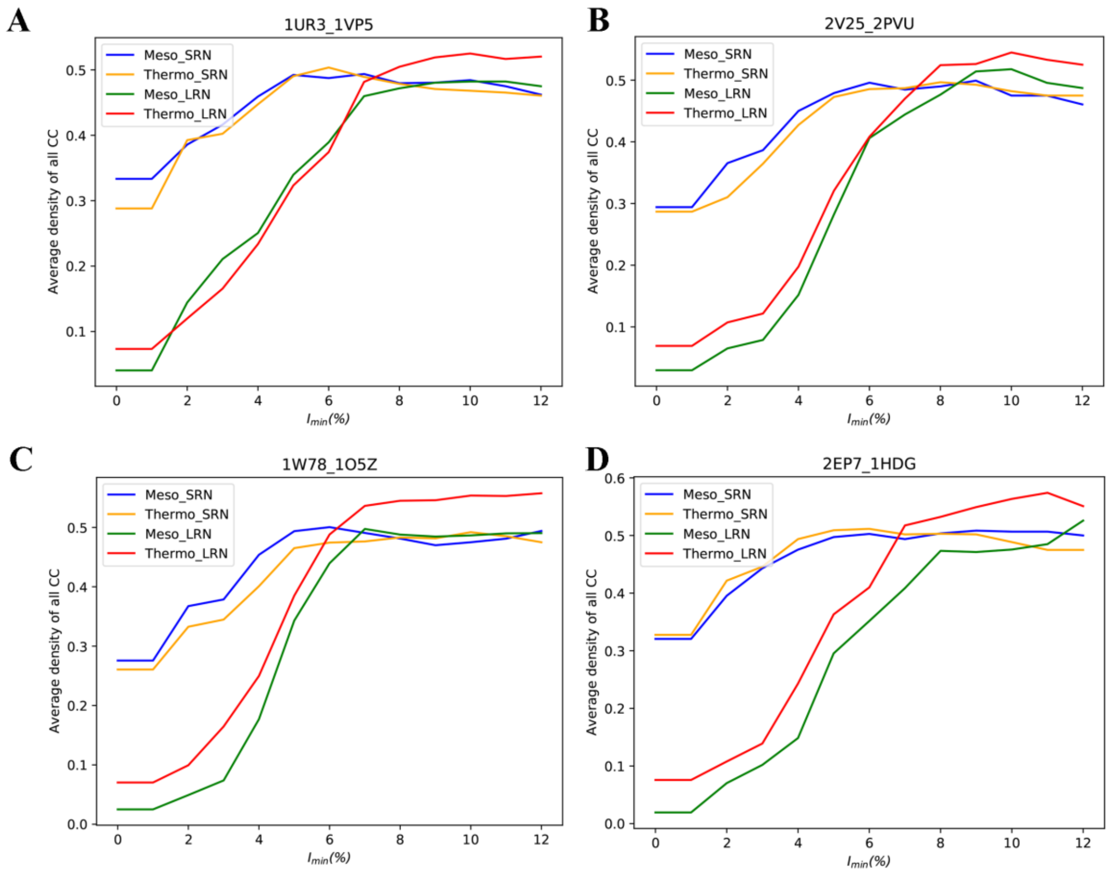


**Figure S1(A-D).** Sample mean ND vs $I_{min}$ plot of four out of 1560 mesophilic-thermophilic orthologous pairs. The mean plot is provided in the main text.

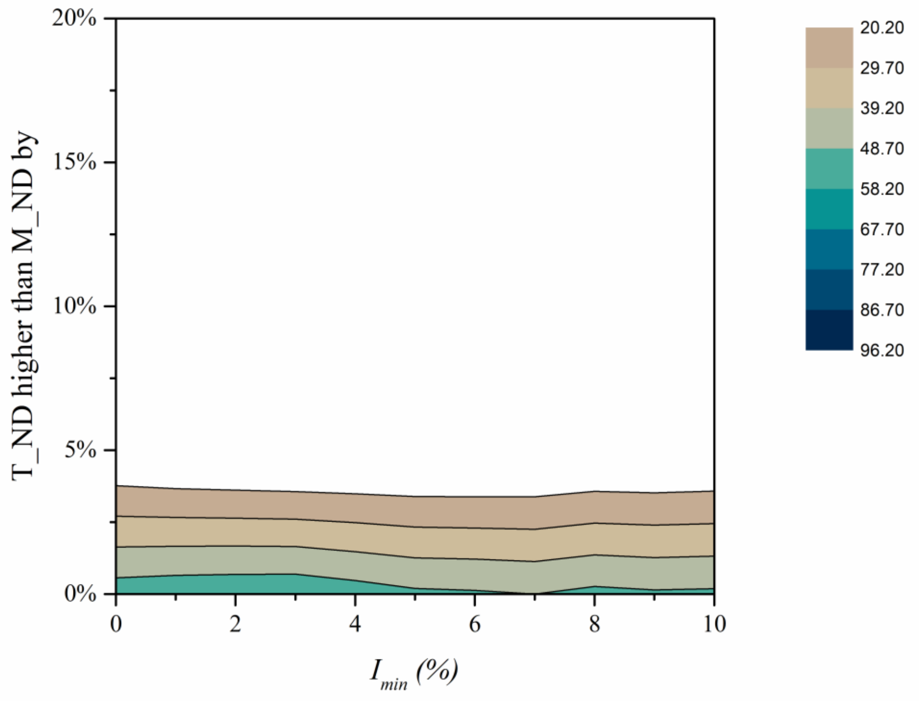


**Figure S3.** The percentages of mesophilic-thermophilic orthologous protein pairs for which average ND values are bigger in thermophilic are plotted as colored contour plots in the the $I_{sel} - \Delta ND_{SRN}$ space.
